# Neuropathology in an α-synuclein preformed fibril mouse model occurs independent of the Parkinson’s disease-linked lysosomal ATP13A2 protein

**DOI:** 10.1101/2024.08.07.607077

**Authors:** Caio M. Massari, Dylan J. Dues, Alexis Bergsma, Kayla Sipple, Darren J. Moore

**Author notes:** Corresponding Author Van Andel Institute, 333 Bostwick Ave NE, Grand Rapids, MI 49503, USA.

## Abstract

Loss-of-function mutations in the *ATP13A2* (*PARK9*) gene are implicated in early-onset autosomal recessive Parkinson’s disease (PD) and other neurodegenerative disorders. *ATP13A2* encodes a lysosomal transmembrane P_5B_-type ATPase that is highly expressed in brain and specifically within the substantia nigra. Recent studies have revealed its normal role as a lysosomal polyamine transporter, although its contribution to PD-related pathology remains unclear. Cellular studies report that ATP13A2 can regulate α-synuclein (α-syn) secretion via exosomes. However, the relationship between ATP13A2 and α-syn in animal models remains inconclusive. *ATP13A2* knockout (KO) mice exhibit lysosomal abnormalities and reactive astrogliosis but do not develop PD-related neuropathology. Studies manipulating α-syn levels in mice lacking ATP13A2 indicate minimal effects on pathology. The delivery of α-syn preformed fibrils (PFFs) into the mouse striatum is a well-defined model to study the development and spread of α-syn pathology. In this study, we unilaterally injected wild-type (WT) and homozygous *ATP13A2* KO mice with mouse α-syn PFFs in the striatum and evaluated mice for neuropathology after 6 months. The distribution, extent and spread of α-syn aggregation in multiple regions of the mouse brain was largely independent of ATP13A2 expression. The loss of nigrostriatal pathway dopaminergic neurons and their nerve terminals induced by PFFs were equivalent in WT and *ATP13A2* KO mice. Reactive astrogliosis was induced equivalently by α-syn PFFs in WT and KO mice but was significantly higher in *ATP13A2* KO mice due to pre-existing gliosis. We did not identify asymmetric motor disturbances, microglial activation, or axonal damage induced by α-syn PFFs in WT or KO mice after 6 months. Although α-syn PFFs induce an increase in lysosomal number in the substantia nigra in general, TH-positive dopaminergic neurons did not exhibit either increased lysosomal area or intensity, regardless of *ATP13A2* genotype. Our study evaluating the spread of α-syn pathology reveals no exacerbation of α-syn pathology, neuronal loss, astrogliosis or motor deficits in *ATP13A2* KO mice, suggesting that selective lysosomal abnormalities resulting from ATP13A2 loss do not play a major role in α-syn clearance or propagation *in vivo*.

## Introduction

Loss-of-function mutations in the *ATP13A2* (*PARK9*) gene cause an early-onset form of autosomal recessive Parkinson’s disease (PD) as well as other related neurodegenerative diseases (Ramirez et al., 2006). *ATP13A2* encodes a lysosomal transmembrane P_5B_-type ATPase that is widely expressed in human tissues, with enriched in the brain and specifically the substantia nigra (Ramirez et al., 2006; Schultheis et al., 2004). Recent studies have elucidated a critical function for ATP13A2 as a lysosomal polyamine exporter (van Veen et al., 2020) although it is still currently unknown how ATP13A2 dysfunction triggers neuropathology in PD. Several cellular studies indicate that ATP13A2 depletion or disease-linked mutations can induce lysosomal dysfunction (Dehay et al., 2012; Matsui et al., 2013; van Veen et al., 2020), and also suggest that ATP13A2 can regulate the extracellular secretion of exosomes containing α-synuclein (α-Syn) (Kong et al., 2014; Tsunemi et al., 2014; Tsunemi & Krainc, 2014). α-Syn fibrils are a principal protein component of Lewy bodies, a pathological hallmark of PD. Interestingly, dopaminergic neurons derived from induced pluripotent stem cells (iPSCs) of patients carrying *ATP13A2* mutations show an accumulation of α-Syn, consistent with PD pathology (Mazzulli et al., 2016; Tsunemi et al., 2020). It has been suggested that *ATP13A2* mutations that result in lysosomal dysfunction could lead to elevated levels of α-Syn that potentially facilitates its aggregation. However, in animal models, the relationship between ATP13A2 and α-Syn in brain cells has not been confirmed.

*ATP13A2* knockout (KO) mice have been developed and well-characterized, and are reported to show widespread age-dependent abnormalities in lysosomal function, the accumulation of lipofuscin, reactive gliosis, and a modest yet selective increase in detergent-insoluble α-Syn in the hippocampus (Kett et al., 2015; Schultheis et al., 2013). However, these KO mice do not exhibit nigrostriatal pathway dopaminergic neurodegeneration or Lewy-like pathology, even with advanced age (∼27 months), indicating that *ATP13A2* KO alone is insufficient to replicate PD-like neuropathology within the mouse lifespan (Kett et al., 2015; Schultheis et al., 2013). PD is a complex disease to model in mammals, likely requiring a combination of genetic and environmental insults combined with aging, in order to trigger robust neurodegeneration. Prior studies have attempted to explore the relationship between α-Syn and ATP13A2 *in vivo*. *ATP13A2* deletion has minimal effects on phenotypes in transgenic mice expressing human wild-type α-Syn, inducing a modest exacerbation of sensorimotor dysfunction (Dirr et al., 2018). Similarly, the overexpression of human ATP13A2 was not sufficient to protect against pathology and neurodegeneration induced by the AAV-mediated expression of human wild-type α-Syn in the substantia nigra of adult rats (Daniel et al., 2015). Key age-related phenotypes in *ATP13A2* KO mice, including histopathology (i.e. gliosis, lipofuscinosis, ubiquitin-protein aggregates, endolysosomal abnormalities) and motor deficits, were shown to occur independently of α-Syn expression (Kett et al., 2015). Moreover, no exacerbation of α-Syn pathology and spread was observed in young heterozygous *ATP13A2* KO mice following the delivery of α-Syn preformed fibrils (PFFs) to the olfactory bulb (Johnson et al., 2021). Since heterozygous *ATP13A2* KO mice are not likely to recapitulate the pathogenic effects of familial recessive mutations, the olfactory PFF study above highlights the importance of investigating the impact of homozygous *ATP13A2* deletion in mice. Furthermore, focusing on brain regions where ATP13A2 is highly expressed and that are most vulnerable in PD, such as the nigrostriatal dopaminergic pathway, is another important consideration.

Accordingly, to further evaluate the impact of ATP13A2 loss-of-function on α-Syn pathology, we unilaterally injected α-Syn PFFs into the striatum of homozygous *ATP13A2* KO mice. Striatal α-Syn PFF delivery provides a useful, well-defined and widely used mouse model of PD to investigate the development of α-Syn pathology and its spread to connected neural circuits (Fares et al., 2016; Henderson et al., 2019; Izco et al., 2021; Luk et al., 2016; Luk et al., 2012; Masuda-Suzukake et al., 2014). We find that the extent and spread of α-Syn pathology throughout the brain, and the resulting nigral dopaminergic neuronal loss, lysosomal accumulation, and reactive astrogliosis, occur independent of ATP13A2 expression. Overall, our study supports the concept that ATP13A2 loss-of-function and associated lysosomal deficits are not sufficient to modulate the clearance or propagation of α-Syn pathology in the nigrostriatal pathway and other neural circuits.

## Materials and Methods

### Animals

*ATP13A2* floxed mice containing *loxP* sites flanking exons 2-3 on a C57BL/6J background were obtained from The Jackson Laboratory (*Atp13a2^tm1.1Wtd^*/J; JAX strain #028387) (Kett et al., 2015). *ATP13A2* floxed mice were crossed with CMV-Cre transgenic mice (B6.C-Tg(CMV-cre)1Cgn/J; JAX strain #006054) to generate germline *ATP13A2* KO progeny, and the CMV-Cre transgene was removed from the resulting KO line via backcrossing to C57BL/6J mice. Cohorts of male and female *ATP13A2* KO mice and their wild-type littermates were developed by interbreeding heterozygous KO mice. Mice were housed and maintained under a 12-h light/dark cycle with free access to food and water at the Vivarium Core of Van Andel Institute. All procedures were performed in accordance with the Guide for the Care and Use of Laboratory Animals (United States National Institutes of Health) and were approved by the Van Andel Institute’s Institutional Animal Care and Use Committee (IACUC).

### Preparation and stereotactic injection of mouse **α**-synuclein PFFs

Recombinant mouse α-synuclein preformed fibrils (PFFs) were produced and validated according to previously established methods (Dues et al., 2023; Volpicelli-Daley et al., 2014; Wang et al., 2019). Fibrils were diluted to a concentration of 5 μg/μl for experimentation. Prior to injection, the fibrils underwent thawing to room temperature and sonication for 4 minutes using a Q700 cup horn sonicator (Qsonica) at 50% power for 120 pulses (1 second on, 1 second off). Male and female 3-4 month-old homozygous *ATP13A2* KO mice and their wild-type littermates were anesthetized with 2% isoflurane and positioned within a stereotactic frame (Stoelting). Mice received a unilateral injection of 10 µg of mouse α-synuclein PFFs in the striatum using the following coordinates in relation to bregma: anterior-posterior (A-P), +1.0 mm; medio-lateral (M-L), +2.0 mm; dorso-ventral (D-V), −2.6 mm. Each injection was delivered in a volume of 2 µl at a flow rate of 0.2 µl/min. All mice were sacrificed at 6 months post-injection (MPI). A total of 14 mice were injected with PFFs. 7 wild-type mice (5 male, 2 female) and 7 homozygous *ATP13A2* KO mice (4 male, 3 female) were utilized. One male *ATP13A2* KO mouse died during recovery from surgery.

### Cylinder test

Forelimb use was evaluated by placing each mouse in a transparent cylinder measuring 15 cm in diameter and 20 cm in height, and their behavior was recorded over a 5-minute period. A camera was positioned above the cylinder to capture forelimb movements from various angles. The contact of either left or right forelimbs with the cylinder wall was measured without knowledge of the experimental groups. The final data was presented as percentage of the ipsilateral (right) forepaw use by calculation with the equation: [(ipsilateral paw + 0.5 both paws) / (ipsilateral paw + contralateral paw + both paws)] x 100. The calculated percentage declares the preference of the forepaw use as follows: 50% = symmetric use of both forepaws; <50% = preference of the contralateral forepaw; >50% = preference of the ipsilateral forepaw (Ip et al., 2017). The test was conducted at 1, 3 and 6 months after PFF injections.

### Immunohistochemistry

Animals were perfused by transcardial perfusion with 0.9% NaCl followed by 4% paraformaldehyde (PFA) in 0.1 M phosphate buffer (PB) at pH 7.4. After perfusion, the whole brain was removed and incubated in 4% PFA in 0.1 M PB at 4°C overnight. Brain tissue was cryoprotected using 30% sucrose in 0.1 M PB for 24 hours before microtome sectioning to prepare 35 µm-thick coronal sections. Brain sections were treated to quench endogenous peroxidase activity by immersing them in 3% H_2_O_2_ (Sigma) diluted in methanol for 5 minutes at 4°C. Subsequently, sections were blocked with 10% normal goat serum (Invitrogen) and 0.1% Triton-X100 in PBS for 1 hour at room temperature. Following this, sections were incubated with primary antibodies for 48 hours at 4°C, followed by biotinylated secondary antibody incubation for 24 hours at 4°C. After incubation with ABC reagent (Vector Labs) for 1 hour at room temperature and visualization using 3, 3’-diaminobenzidine tetrahydrochloride (DAB; Vector Labs), sections were mounted on Superfrost Plus slides (Fisher Scientific). Slide-mounted sections were then dehydrated using increasing concentrations of ethanol and xylene, before being coverslipped with Entellan (Merck). Primary antibodies against pS129-α-synuclein (ab51253; Abcam), tyrosine hydroxylase (TH; N300-109; Novus Biological), Iba1 (019-19741; Fujifilm Wako Chemical USA), and GFAP (ab227761; Abcam) were used. Biotinylated goat anti-rabbit and goat anti-mouse (Vector labs) secondary antibodies were used.

### Immunofluorescence

Brain sections were incubated with primary antibodies, chicken anti-TH (ab76442; Abcam) and rat anti-LAMP2 (ab13524; Abcam), at 4°C overnight followed by incubation with secondary antibodies goat anti-rat AlexaFluor-546 (A-11081), and goat anti-chicken AlexaFluor-647 (A-21449) from Thermofisher Scientific and DAPI for nuclei staining. Sections were mounted on Superfrost Plus slides (Fisher Scientific) and confocal images were captured with an ImageXpress® Confocal HT.ai High-Content Imaging System using a 60x air objective and analyzed using IN Carta® Image Analysis Software (Molecular Devices).

### Digital and quantitative pathology

Slide-mounted coronal brain sections were scanned using a ScanScope XT slide scanner (Aperio) at a resolution of 0.5 μm/pixel. Annotation and quantitative assessment of the brain regions of interest (forebrain, cortex, striatum, amygdala, and substantia nigra) were performed using HALO analysis software (Area quantification and Microglial activation modules; Indica Labs Inc.). Contralateral and ipsilateral (relative to the PFF injection site) hemispheres were annotated and assessed separately. For each quantitative approach, threshold parameters were developed and optimized to detect positive-staining but excluding background staining or tissue-level artifacts. All sections/slides were assessed using the optimized HALO batch analysis function for each marker of interest. All annotations and assessments were conducted by researchers blinded to slide identification or animal genotype.

### Optical density analysis of striatal TH-positive terminals

DAB immunostaining was used to label TH in coronal brain sections containing the striatum. Images were obtained using an Aperio ScanScope XT slide scanner. Mean optical density in the striatum was measured in every 4^th^ section using HALO analysis software (Area quantification module; Indica Labs Inc.).

### Stereological quantification of substantia nigra TH-positive neurons

Ventral midbrain sections were immunostained for TH and counterstained with 0.1% cresyl violet. TH-positive and total Nissl-positive neurons in the substantia nigra were estimated using unbiased stereological quantification. Every 4^th^ serial section of the ventral midbrain was evaluated using the optical fractionator probe of the StereoInvestigator software (MicroBrightField Biosciences). Random, systematic sampling was performed using a grid of 120 x 120 µm squares and applying an optical dissector with the dimensions 50 x 50 x 14 µm. During analysis, investigators were blinded to experimental conditions.

### Gallyas silver staining

To analyze axonal damage, we performed Gallyas silver staining using the NeuroSilver^TM^ Kit II (FD NeuroTechnologies Inc) according to the manufacturer’s instructions.

### Statistical analysis

Data was analyzed with GraphPad Prism 9 software by two-tailed paired or unpaired Student’s *t*-test, one-way or two-way ANOVA with Tukey’s multiple comparisons test as described in each figure legend. Graphs were generated using GraphPad Prism 9 and depict all data as mean ± SEM.

## Results

### Delivery of α-Syn PFFs into the mouse striatum of WT and *ATP13A2* KO mice

For this study, our experimental design consisted of a unilateral injection of 10 ug of recombinant mouse α-Syn PFFs into the striatum of KO mice lacking the lysosomal protein ATP13A2 or their wild-type (WT) littermates and subsequent pathological investigation at 6 months post-injection (MPI) (**Figure 1A**). To evaluate motor disturbance, 30, 90 and 180 days after the α-Syn PFF injection, mice were subjected to the cylinder test. A decrease in the total number of forelimb contacts on the cylinder wall were seen at 6 MPI but both contralateral and ipsilateral paw touches remained at approximately 50%, therefore no paw preference was observed, indicating that α-Syn PFF injection in the striatum in WT mice did not cause asymmetric motor behavior (**Figure 1B-C**). Of note, mice lacking ATP13A2 also did not present asymmetric motor preferences, while a slight decrease in total paw touches was already seen at 3 MPI (**Figure 1B-C**).

**Figure 1.**
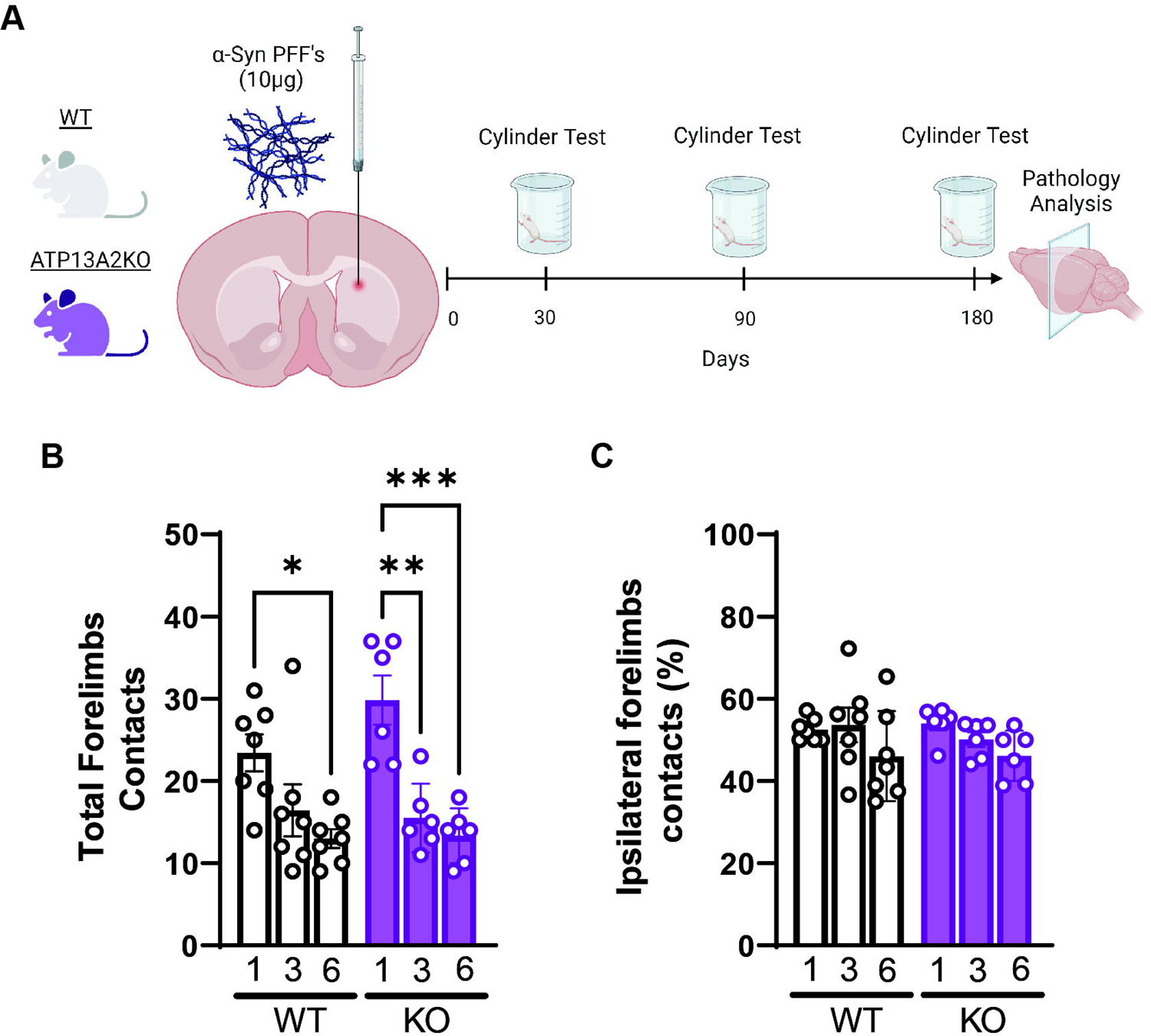
Schematic illustration of experimental design and forelimb use. **(A)** Adult WT and *ATP13A2* KO mice were subjected to unilateral stereotactic injection of α-synuclein PFFs (10 µg) into the striatum. Cohorts were assessed at 1, 3 and 6 MPI for motor asymmetry by cylinder test and at 6 MPI for neuropathological analyses. Total (B) and % ipsilateral (**C**) forelimb contacts with the cylinder wall were counted over 5 min at 1, 3 and 6 MPI. Each circle represents one animal, and bars depict mean ± SEM (*n* = 7 for WT and *n* = 6 for *ATP13A2* KO). (**C**) The calculated percentage indicates forepaw use preference as follows: 50% = symmetric use of both forepaws; <50% = preference for the contralateral forepaw; >50% preference for the ipsilateral forepaw. \**P*<0.05 or \*\**P*<0.01 by two-way ANOVA with Tukey post-hoc analysis.

### Pathological **α**-Syn distribution is not altered by *ATP13A2* KO

Previous studies have already mapped, in detail, pathological α-Syn distribution and burden in similar striatal PFF models (Henderson et al., 2019; Luk et al., 2012; Masuda-Suzukake et al., 2014). Consistent with prior studies, we observe that pathologic α-Syn accumulation occurs throughout the mouse brain. We focused on the regions with the most prominent α-Syn burden, namely, the forebrain, cortex, striatum, amygdala and substantia nigra pars compacta (SNpc) (**Figure 2A, D, G, J, M**). Overall, in all these brain regions we observe a clear increase of pathological α-Syn staining, measured by immunoreactivity for pS129-α-Syn (**Figure 2B, E, H, K, N**). While in some brain regions pS129-α-Syn is detected in both hemispheres, we observe a significant increase for pS129-α-Syn staining in the injected hemisphere compared to the non-injected hemisphere in all brain regions analyzed. The same pattern is seen in both WT and *ATP13A2* KO animals, showing no significant differences between genotypes related to propagation or accumulation of pS129-α-Syn in the brain. To evaluate the level of α-Syn increase between hemispheres, the ratio of pathological α-Syn in the ipsilateral versus the contralateral injection side was also measured, with no difference among genotypes (**Figure 2C, F, I, L, O**). Our data provides evidence that ATP13A2 may not play a direct role in the clearance of α-Syn accumulation or its propagation *in vivo*.

**Figure 2.**
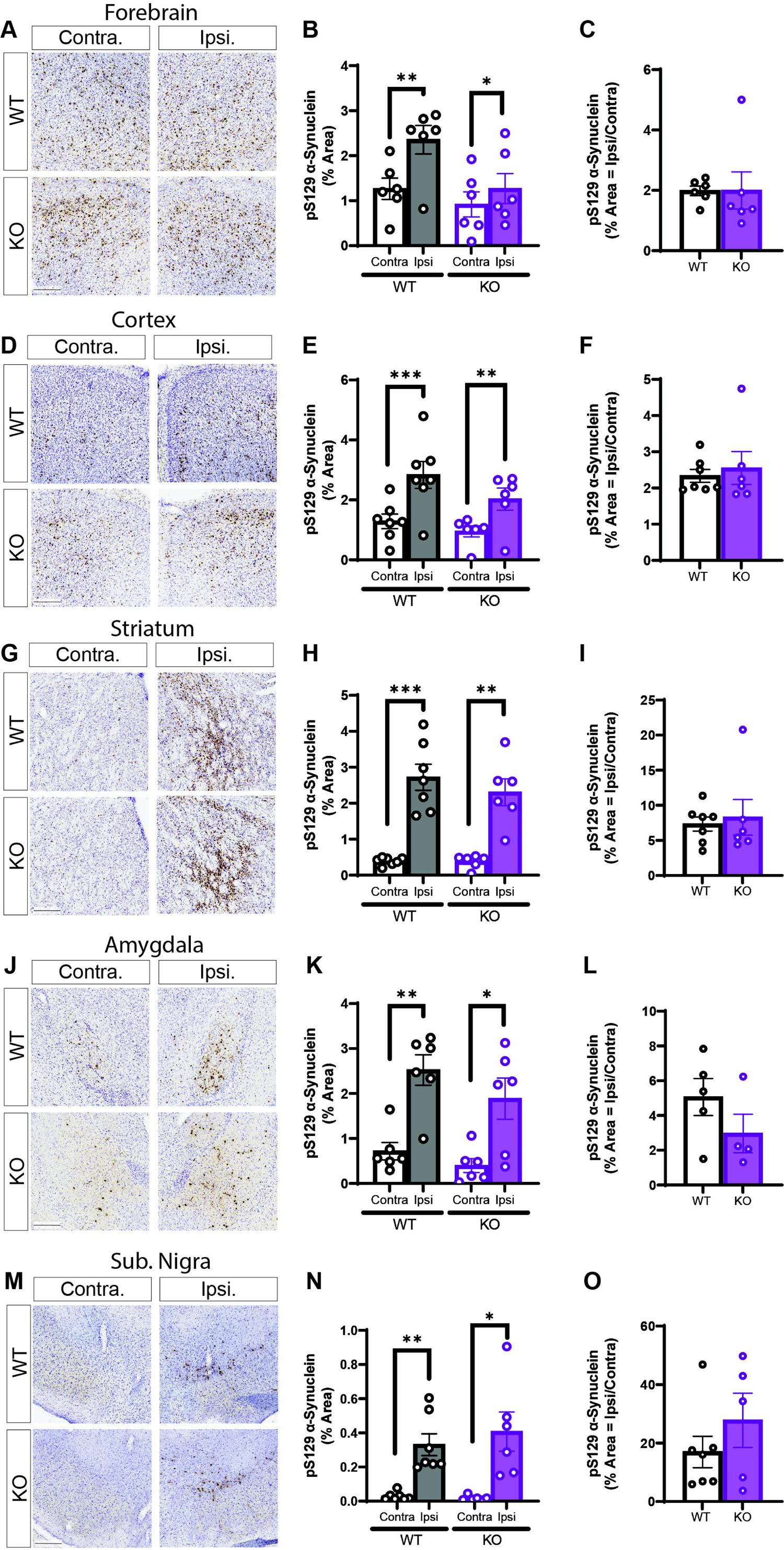
No impact of *ATP13A2* KO on α-synuclein pathology in the mouse brain at 6 MPI. (**A**, **D**, **G**, **J**, and **M**) Representative images of pS129-α-synuclein immunostaining at 6 MPI in WT or *ATP13A2* KO mice, from the contralateral and ipsilateral (injected) hemispheres. Scale bar: 200 µm. Digital pathology quantitation of pS129-α-synuclein immunostaining in the (**B**) forebrain, (**E**) cortex, (**H**) striatum, (**K**) amygdala, and (**N**) substantia nigra at 6 MPI. Data are expressed as the % area occupied by pS129-α-synuclein immunostaining in the contralateral or ipsilateral hemisphere, with bars representing the mean ± SEM, and individual data points for each animal (*n* = 6–7 animals/group). \**P*<0.05 or \*\**P*<0.01 by paired Student’s *t*-test comparing contralateral and ipsilateral hemispheres within the same genotype as indicated, and no significance between genotypes by two-way ANOVA with Tukey’s multiple comparisons test. (**C**, **F**, **I**, **L**, and **O**) Ipsilateral to contralateral ratio of % area of pS129-α-synuclein immunostaining for each animal within each brain region, with bars representing mean ± SEM for each genotype. *P*>0.05 by unpaired Student’s *t*-test.

### TH-positive neuronal loss induced by **α**-Syn PFFs is not aggravated in *ATP13A2* **KO mice**

One of the key effects of α-Syn PFF injection into the mouse striatum is the degeneration of substantia nigra dopaminergic neurons, highlighting the relevance of this model to PD. Previous studies have already shown that α-Syn PFF injection in the striatum leads to striatal TH-positive nerve terminal loss and SNpc TH-positive cell loss beginning by 90 days post-injection (Henderson et al., 2019; Luk et al., 2016; Luk et al., 2012). Here, we analyzed the density of TH-positive staining in the striatum and estimated neuronal cell number in the SNpc by unbiased stereology at 180 days after PFF injection. When comparing the α-Syn PFF-injected hemisphere to the non-injected contralateral hemisphere, we find ∼40% loss of TH fiber density in the striatum and ∼50% loss of TH-positive cells in the SNpc due to α-Syn PFFs (**Figure 3B, F**). While a robust PFF-induced lesion is observed in both genotypes, we do not find significant differences in TH-positive lesion size between WT and *ATP13A2* KO mice (**Figure 3C, G**). These data indicate that the lack of ATP13A2 does not further sensitize dopaminergic neurons to the pathological effects of α-Syn accumulation.

**Figure 3.**
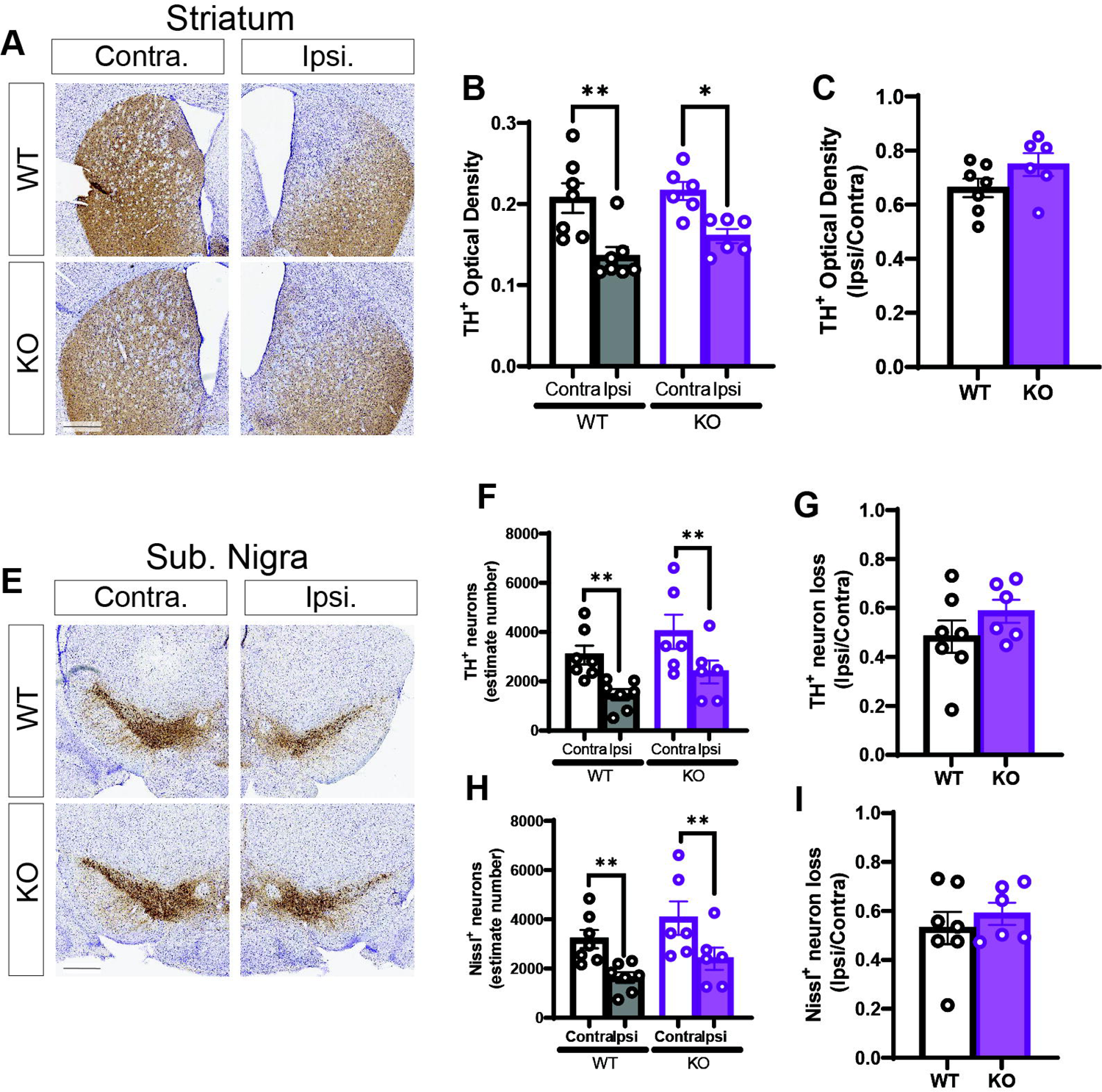
Nigrostriatal TH-positive dopaminergic cell and terminal loss induced by α-synuclein PFFs is not altered by *ATP13A2* KO. Representative images of tyrosine hydroxylase (TH) immunostaining in the striatum (**A**) and substantia nigra pars compacta (**E**) of WT or KO mice at 6 MPI of α-synuclein PFFs. Scale bars: 500 µm. (**B**) Striatal TH-positive optical density in the ipsilateral and contralateral hemispheres of WT or KO mice was measured using HALO analysis software. Bars represent mean ± SEM, and circles represent individual animals (*n* = 6-7 mice/genotype). \**P*<0.05 or \*\**P*<0.01 by paired Student’s *t*-test comparing contralateral and ipsilateral TH-positive optical density within each genotype as indicated, with no significance comparing TH-positive optical densities between genotypes by two-way ANOVA with Tukey’s multiple comparisons test. (**C**) Ipsilateral to contralateral ratio of striatal TH-positive density for each genotype, with no significance by unpaired, Student’s *t*-test. Bars represent mean ± SEM, and circles represent individual animals (*n* = 6-7 mice/genotype). Unbiased stereological analysis of TH-positive dopaminergic (**F**) and total Nissl-positive (**H**) neurons in the substantia nigra of ipsilateral and contralateral hemispheres from WT and KO mice, with data expressed as estimates of TH-positive or Nissl-positive neuron numbers (mean ± SEM, *n* = 6-7 mice/group). \*\**P*<0.01 comparing contralateral and ipsilateral hemispheres within each genotype by paired Student’s *t*-test, and no significance comparing cell numbers between genotypes by two-way ANOVA with Tukey’s multiple comparisons test. (**G**, **I**) Ipsilateral to contralateral ratio of nigral TH-positive or Nissl-positive neuron loss for WT and KO mouse genotypes, with no significance by unpaired, Student’s *t*-test.

### Lack of striatal axonal degeneration induced by PFFs

In order to detect active neurite degeneration rather than the loss of nerve terminals, we used Gallyas silver staining in our α-Syn PFF model. Previous studies have shown that α-Syn can induce changes in spine density, dynamics, and morphology that lead to spine loss in neurons (Blumenstock et al., 2017; Froula et al., 2018; Volpicelli-Daley et al., 2014). In our hands, silver-positive degenerating neuronal processes are not robustly induced by α-Syn PFFs at 6 months. We find a rather modest increase of axonal degeneration in the ipsilateral striatum (**Figure 4A**) and minimal degeneration in the ipsilateral SNpc of WT mice (**Figure 4B**). However, the burden of neurite degeneration in ipsilateral hemispheres of WT and *ATP13A2* KO mice is indistinguishable. Overall, we find no evidence for an α-Syn PFF-induced aggravation of axonal degeneration in the absence of *ATP13A2*.

**Figure 4.**
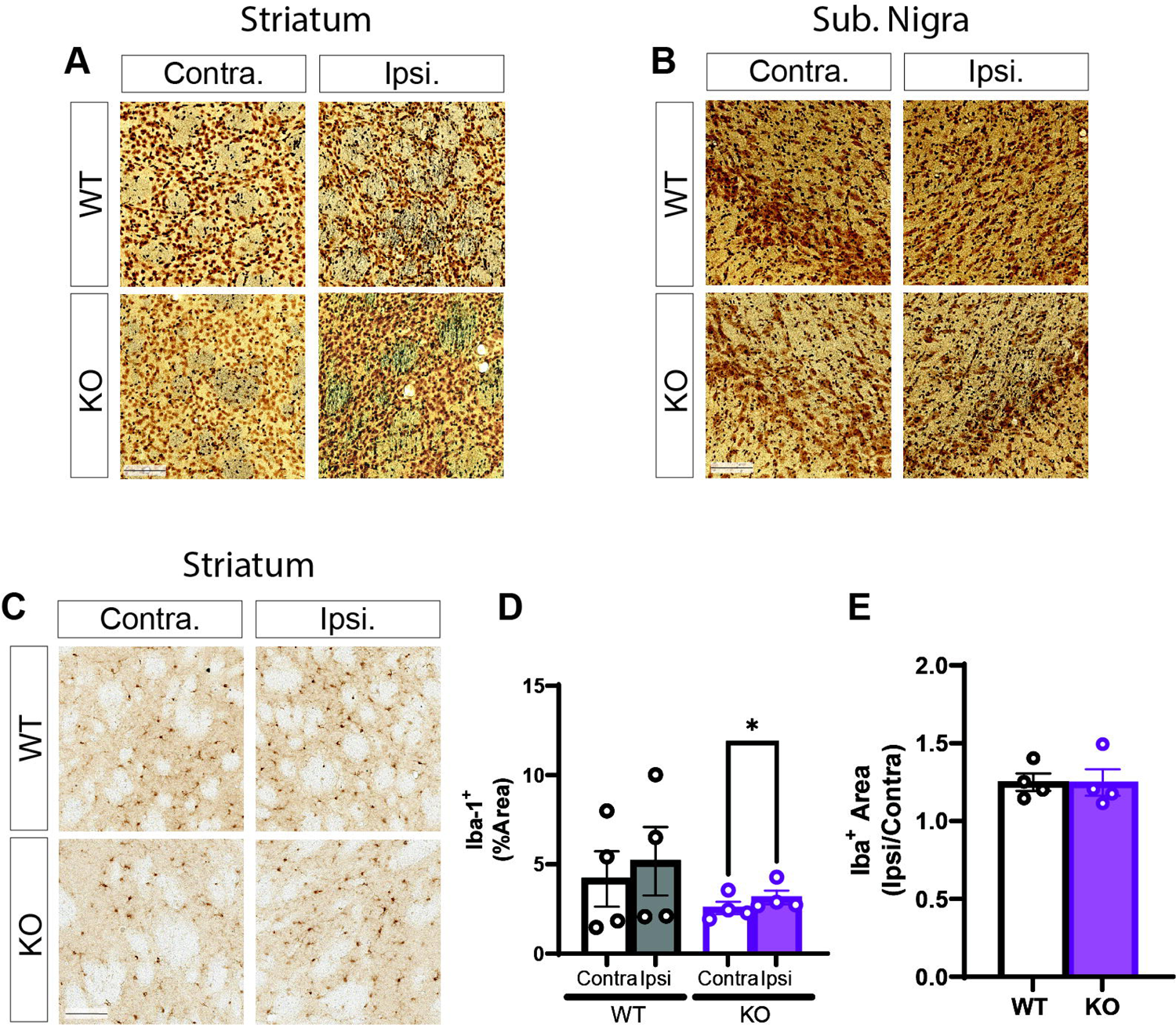
Lack of axonal degeneration or microglial activation induced by α-synuclein PFFs and *ATP13A2* deletion. Representative images of Gallyas silver staining (gray/black fibers or puncta) from striatum (**A**) and substantia nigra (**B**) of WT and KO mice, comparing contralateral with ipsilateral PFF injection at 6 MPI. Data are representative of *n* = 4 animals/genotype. (**C**) Representative images of ipsilateral (PFFs) and contralateral striatum from WT or KO mice immunostained for Iba-1 to detect microglia. (**D**) Data are expressed as % area of Iba-1 immunostaining with bars representing mean ± SEM, and circles indicating individual animals (*n* = 4 mice/group). \**P*<0.05 by paired Student’s *t*-test comparing contralateral and ipsilateral % Iba-1 area within each genotype, with no significance between genotypes by two-way ANOVA with Tukey’s multiple comparisons test. (**E**) Ipsilateral to contralateral ratio of % Iba-1 area for WT and KO mice indicates a modest increase in microglial density, but with no significance between genotypes by unpaired, Student’s *t*-test. (**A-C**) Scale bars: 100 µm.

### Reactive astrogliosis induced by **α**-Syn PFFs or *ATP13A2* deletion

Since inflammation plays an important role in the neurodegenerative process we sought to determine if microglia and astrocytes are altered by α-Syn PFFs and the impact of the absence of ATP13A2. Microglial activation presents as an early response to α-Syn PFF delivery but little is known beyond 90 days post-injection (Izco et al., 2021). In our model, at 6 MPI, we do not observe robust microglial changes due to α-Syn PFF delivery. Cortex, amygdala and SNpc do not present an increase in Iba-1 immunostaining in the ipsilateral hemisphere or pronounced changes in morphology consistent with their activation (**Figure S1**). Iba1-positive immunostaining in the striatum is modestly increased in the ipsilateral compared to the contralateral hemisphere of *ATP13A2* KO mice (**Figure 4D**), but this effect is not observed in WT mice. However, when we compare the Iba-1 immunostaining ratio of ipsilateral over contralateral hemispheres among genotypes there is no significant difference, indicating that the level of microglial density is comparable in these animals and that the small increase is likely not biologically relevant (**Figure 4E**).

For astrocytes, immunostaining for GFAP reveals a robust increase of this astrocytic marker in the α-Syn PFF-injected hemisphere, mainly in cortex and striatum (**Figure 5A, D**). The amygdala and SNpc do not display a significant increase in GFAP levels (**Figure S2**). Of note, *ATP13A2* KO mice show significantly higher levels of GFAP staining in the cortex and striatum, both in the injected and non-injected hemispheres, when compared to WT animals (**Figure 5B-C, E-F**). This increase was expected, since the initial characterization of *ATP13A2* KO mice reported widespread astrogliosis as an important age-related phenotype of these mice (Kett et al., 2015). Interestingly, the GFAP immunostaining ratio of ipsilateral over contralateral hemispheres between genotypes are not different for cortex but are significantly reduced for striatum in KO mice (**Figure 5C, F**). It is plausible that *ATP13A2* KO mice already have near saturated levels of reactive astrogliosis and exhibit a smaller response when exposed to α-Syn PFF-induced pathology, relative to their robust effects in WT mice. Our data suggests that astrocytes might be more affected by the lack of ATP13A2 than other brain cell types.

**Figure 5.**
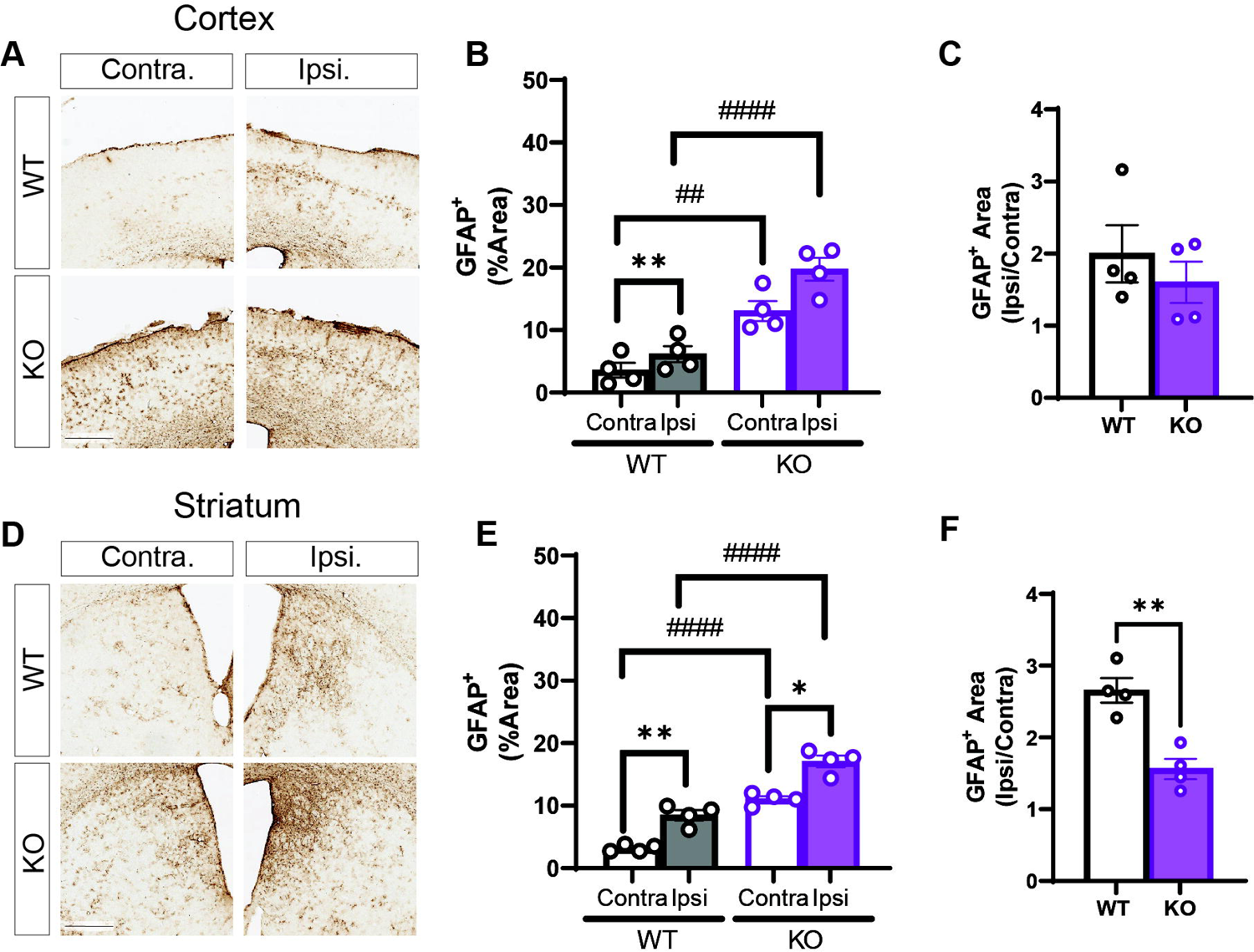
Reactive astrogliosis induced by *ATP13A2* KO or α-synuclein PFF delivery. Representative images for GFAP immunostaining from cortex (**A**) and striatum (**D**) of WT and KO mice, comparing contralateral with ipsilateral PFF injection at 6 MPI. Scale bars: 500 µm. (**B, E**) Digital pathology quantitation of GFAP immunostaining in (**B**) cortex and (**E**) striatum expressed as % area, with bars representing mean ± SEM and circles representing individual animals (*n* = 4 animals/group). \**P*<0.05 and \*\**P*<0.01 by paired Student’s *t*-test comparing contralateral and ipsilateral hemispheres within each genotype, and ^##^*P*<0.01 and ^####^*P*<0.0001 by two-way ANOVA with Tukey’s multiple comparisons test comparing between genotypes. (**C, F**) Ipsilateral to contralateral ratio of % GFAP area for WT and KO mice in (**C**) cortex and (**F**) striatum, indicates a robust increase in astrocyte density induced by PFFs, that is significantly reduced in the striatum of KO mice. \*\**P*< 0.01 by unpaired Student’s *t*-test.

### Lysosomal accumulation in the substantia nigra induced by **α**-Syn PFFs

Since ATP13A2 is a lysosomal transmembrane protein, we sought to evaluate if SNpc dopaminergic neurons would develop lysosomal damage. Using IN Carta® Image Analysis software, we created a mask and digitally analyzed LAMP2-positive lysosomes in the SNpc and in TH-positive dopaminergic neurons (**Figure 6A, D, G**). As expected, we reproduce a modest decrease in TH-positive neuron number in the SNpc of the ipsilateral hemisphere (**Figure 6B**) with no difference in the TH-positive neuron area among genotypes or hemispheres (**Figure 6C**). Applying the LAMP2 mask to the entire SNpc, we observe an overall increase in LAMP2-positive lysosome counts induced by α-Syn PFFs at 6 MPI (**Figure 6E**) but mean area analysis indicates that these lysosomes are not enlarged or swollen (**Figure 6F**). Finally, the analysis of LAMP2-positive lysosomes in TH-positive neurons reveals no difference in LAMP2 area or fluorescent intensity, regardless of PFF exposure or genotype (**Figure 6H, I**). These data suggest that surviving TH-positive dopaminergic neurons do not exhibit lysosomal damage at 180 days following α-Syn PFF delivery yet indicate that surrounding SNpc non-dopaminergic cells do exhibit lysosomal accumulation.

**Figure 6.**
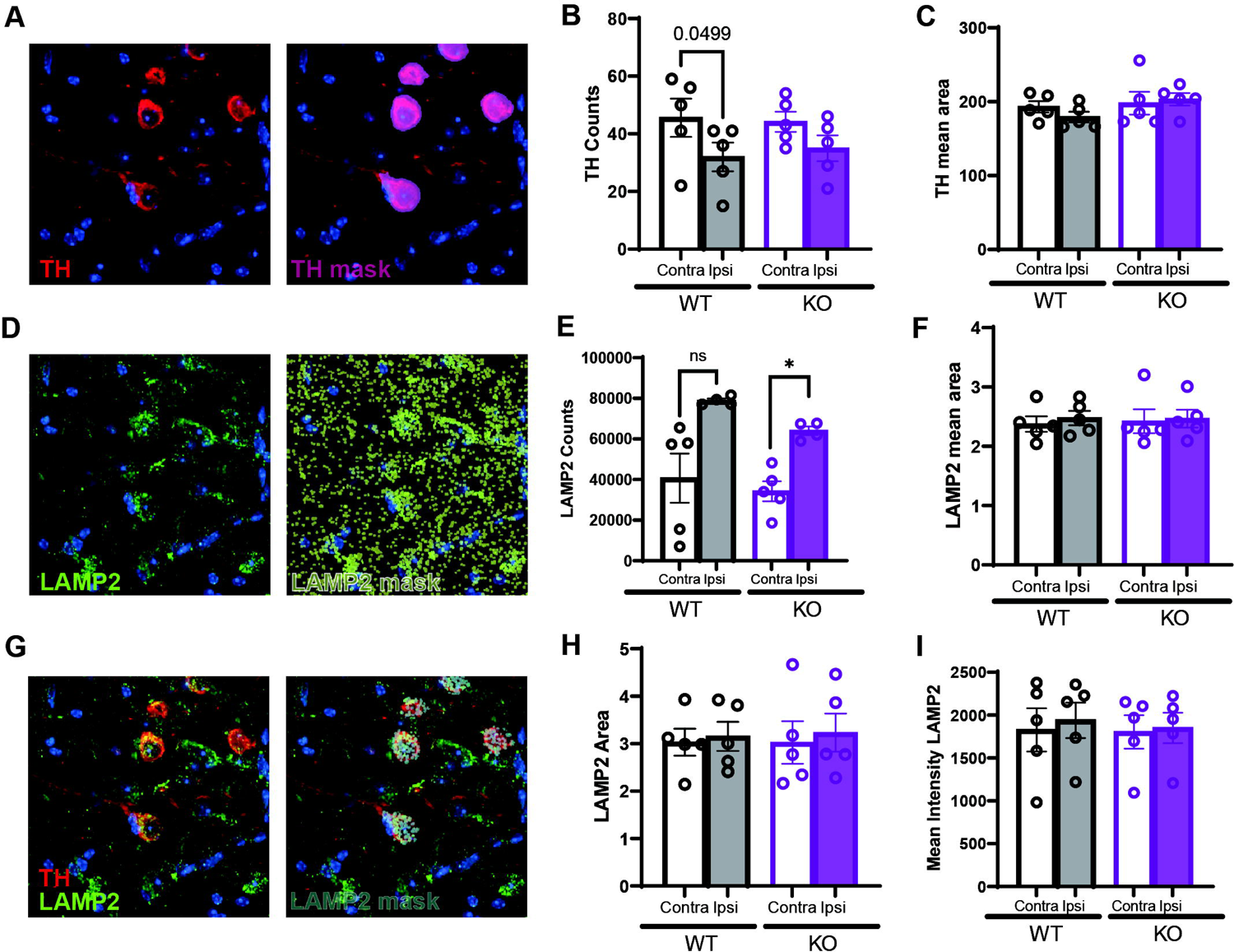
Accumulation of lysosomes in the substantia nigra induced by α-synuclein PFF delivery. Representative immunofluorescent images of TH-positive neurons in the substantia nigra of mice injected with α-synuclein PFFs at 6 MPI. TH-positive immunofluorescence (red) and the corresponding TH-positive mask developed in INCarta (magenta) are shown (**A**). Quantitation of number (**B**) and mean area (**C**) of immunofluorescent TH-positive neurons in ipsilateral (PFFs) or contralateral hemispheres of WT and KO mice, with bars representing mean ± SEM and circles representing individual animals. (**D**) Images of LAMP2-positive immunofluorescence (green) and the corresponding LAMP2-positive mask (light yellow) in the substantia nigra are shown. Quantitation of number and mean area of LAMP2-positive puncta in the substantia nigra in ipsilateral (PFFs) or contralateral hemispheres of WT and KO mice are shown in (**E**) and (**F**), respectively, with bars representing mean ± SEM and circles representing individual animals. (**G**) Colocalization of TH (red) and LAMP2 (green) immunofluorescence are shown, and the corresponding TH-positive LAMP2 mask (cyan). Quantitation of LAMP2-positive mean area (**H**) and fluorescence intensity (**I**) specifically in nigral TH-positive neurons in ipsilateral (PFFs) or contralateral hemispheres of WT and KO mice, with bars representing mean ± SEM and circles representing individual animals. Images were captured at 60x magnification on a confocal microscope and indicate a single z-plane at 0.15 mm thickness. \**P*<0.05 by paired Student’s *t*-test comparing contralateral and ipsilateral hemispheres within each genotype (*n* = 5 mice/group). No differences were found between genotypes by two-way ANOVA with Tukey’s post-hoc test.

## Discussion

The major findings of this study are that α-Syn pathology and neurodegeneration induced by α-Syn PFF delivery in the mouse striatum occurs independent of ATP13A2 expression. While *ATP13A2* loss-of-function mutations cause an early-onset form of autosomal recessive PD, and related neurodegenerative diseases, the precise mechanisms that lead to disease in humans are not well established. Here, we sought to challenge homozygous *ATP13A2* KO mice with α-Syn PFFs to evaluate if the loss of this neuroprotective lysosomal protein would exacerbate PD-like neuropathology *in vivo*. When compared to WT mice, *ATP13A2* KO mice did not reveal significant differences in pathological α-Syn levels and distribution throughout connected neural circuits, nigrostriatal pathway dopaminergic neurodegeneration, neuroinflammatory response, or substantia nigra lysosomal abnormalities. Taken together, *ATP13A2* deletion did not exacerbate or sensitize dopaminergic neurons to the neurotoxic effects of α-Syn PFFs. These findings provide important insight for understanding the role of ATP13A2 in the molecular pathogenesis of PD and its potential relationship with α-Syn *in vivo*.

Since the first description of polyamines as substrates of lysosomal export by ATP13A2 (De La Hera et al., 2013; van Veen et al., 2020), it remains unclear how or if polyamine dysregulation could contribute to PD. Polyamines are a class of widely distributed small, positively-charged alkylamines, which include spermine, spermidine, and their precursor putrescine, and they serve numerous functions, such as cell growth, survival, and proliferation (Pegg, 2016). Polyamines can act as scavengers of free radicals to reduce oxidative stress (Ha et al., 1998; Rider et al., 2007; Sagar et al., 2021). Lysosomal polyamine export mediated by ATP13A2 can lower mitochondrial-derived reactive oxygen species and improve mitochondrial function (Vrijsen et al., 2020). The effects of polyamines may also be directly related to α-Syn. *In vitro* studies, monitored by nuclear magnetic resonance and circular dichroism spectroscopy, showed that putrescine, spermidine, and spermine, at physiological concentrations, can accelerate the aggregation and fibrillization of α-Syn (Antony et al., 2003; Fernandez et al., 2004). This may suggest that polyamine dysregulation in the cytosol due to ATP13A2 loss could potentially support α-Syn aggregation. We speculated that the loss of ATP13A2 and the subsequent polyamine imbalance might aggravate α-Syn neuropathology in mice. Although this was not found in our PFF α-Syn mouse model or in other rodent models expressing human α-Syn (Daniel et al., 2015; Kett et al., 2015), there is some evidence for an interaction in human cellular models. Cultured human fibroblasts derived from subjects harboring *ATP13A2* mutations displayed increased levels of detergent-insoluble α-Syn and decreased levels of secreted or exosomal α-Syn, suggesting that ATP13A2 could play a normal role in the secretion of α-Syn (Tsunemi et al., 2014). A similar effect was later observed in human iPSC-derived dopaminergic neurons obtained from subjects carrying *ATP13A2* mutations (Mazzulli et al., 2016; Tsunemi et al., 2019). Nevertheless, it is still not clear whether patient brains harboring *ATP13A2* mutations present with Lewy body pathology as post-mortem analysis is limited. To our knowledge, only one study has described the neuropathological analysis of a 38-year-old subject with early-onset PD due to homozygous missense mutations in *ATP13A2*, but without detection of Lewy body-like inclusions (Chien et al., 2021).

The α-Syn PFF model can often present with some variability among different laboratories mainly due to differences in PFF preparations (for review see (Polinski, 2021)). Here, we recapitulated the majority of reported pathological features induced by PFFs. We observed a pronounced increase in pS129-α-Syn-positive pathology, not only in the injected region, but also the forebrain, ventral midbrain and amygdala, the robust loss of SNpc dopaminergic neurons and their striatal nerve terminals, and reactive astrogliosis. Of note, delivery of α-Syn PFFs into the mouse striatum produces differences in pathology over time and within brain regions. While microglial activation in the SNpc has been detected in the first 15 days post-injection, it is not present in the striatum or cortex (Izco et al., 2021). Astrogliosis develops later in the striatum and cortex (i.e. 90 days post-injection), but for the SNpc it is evident at 15 days post-injection but absent by 30 days (Izco et al., 2021). Neuroinflammation therefore appears to be an early and transient event in the PFF model. At 180 days post-injection, we no longer find evidence for microglial activation, despite a small increase in Iba-1-positive area in the injected striatum of *ATP13A2* KO mice, although the ratio of ipsilateral over contralateral microglia is similar when compared to WT mice. α-Syn PFF delivery into the olfactory bulb also did not activate microglia after 180 days (Johnson et al., 2021). Astrogliosis induced by α-Syn PFF delivery was robust in the cortex and striatum of WT mice, with a significant further increase in *ATP13A2* KO mice. However, it is known that *ATP13A2* KO mice exhibit progressive astrogliosis in most brain regions (Kett et al., 2015), and the combined effects of PFFs and *ATP13A2* KO appeared to be additive rather than synergistic as suggested by similar increased ratios of ipsilateral/contralateral GFAP-positive astrocytes in WT versus KO mice.

ATP13A2 is a lysosomal protein with high expression in the SNpc (Ramirez et al., 2006; Ramonet et al., 2012). *ATP13A2* KO mice exhibit defective processing of cathepsin D, a lysosomal protease, and increased LAMP1 and LAMP2 expression in the cortex and cerebellum, however, these effects were not present in SNpc dopaminergic neurons (Kett et al., 2015). In our study, we did not observe an increase in LAMP2-positive lysosomal area in TH-positive SNpc neurons of *ATP13A2* KO mice or induced by striatal delivery of α-Syn PFFs. Lysosomal abnormalities are age-dependent in *ATP13A2* KO mice (Kett et al., 2015), and it is possible that the KO mice used here are not aged sufficiently to reveal this phenotype (i.e. 9-10 months). At 6 MPI, since there is already a marked decrease in TH-positive neuron number in the SNpc, these neurons infrequently display pS129-α-Syn-positive inclusions. Our data therefore only represents the lysosomal status of surviving dopaminergic neurons, and it is possible these neurons either were not originally exposed to α-Syn-induced toxicity or were resilient to it. In the SNpc, we did observe an increase in lysosomal number induced by α-Syn PFFs outside of TH-positive neurons, suggesting that other cell types may experience lysosomal stress. Since pS129-α-Syn-positive pathology is restricted to nigral dopaminergic neurons, following striatal PFF delivery, is it possible that reactive astrocytes in the substantia nigra exhibit altered lysosomal number in response to ongoing neuronal cell death.

One caveat of our study is the use of germline *ATP13A2* KO mice, with the potential for developmental compensation that restricts neurodegeneration and their sensitivity to α-Syn PFF delivery. While potential compensatory effects are difficult to ascertain, further investigations are warranted to clarify the relationship between ATP13A2 with α-Syn, particularly given the reported differences between mouse models and human cellular models. To overcome developmental compensation, we recently developed a mouse model with the adult-onset deletion of *ATP13A2*, that now develops progressive dopaminergic neurodegeneration and lysosomal abnormalities in the SNpc, yet without evidence for α-Syn aggregation (Erb et al., 2024). It will be of interest to explore the susceptibility of this new conditional *ATP13A2* KO model to α-Syn PFF exposure. The postmortem analysis of brains from additional *ATP13A2*-linked PD subjects is now needed to further clarify the relationship between ATP13A2 loss-of-function and Lewy body pathology.

## Conclusions

Our study evaluating the extent and spread of α-Syn pathology induced by striatal PFF delivery reveals no exacerbation of α-Syn pathology, neuronal loss, reactive astrogliosis or lysosomal accumulation in *ATP13A2* KO mice, suggesting that lysosomal ATP13A2 does not play a major role in α-Syn clearance or propagation *in vivo*.

## Acknowledgements

We are grateful for funding support from a Pathway-to-Independence (P2i) post-doctoral fellowship awarded by the West Michigan Neurodegenerative Diseases (MiND) Program at VAI (to C.M.M.) and from Van Andel Institute. We thank the Van Andel Institute Pathology and Biorepository Core (RRID:SCR_022912) and Optical Imaging Core (RRID:SCR_021968) (Grand Rapids, MI) for technical assistance.

## Supplementary Figures

**Figure S1. Lack of microglial activation induced by α-synuclein PFFs or *ATP13A2* deletion in distinct brain regions.** Representative images of Iba-1 immunostaining to detect microglia from cortex (**A**), amygdala (**D**) and substantia nigra (**G**) from WT or *ATP13A2* KO mice, comparing contralateral with ipsilateral PFF injection at 6 MPI. Scale bars: 100 µm. (**B, E, F**) Quantitation of Iba-1 immunostaining in each brain region with data expressed as % Iba-1-positive area, with bars representing mean ± SEM and circles indicating individual animals (*n* = 4 mice/group). *P>*0.05 by paired Student’s *t*-test comparing contralateral and ipsilateral % Iba-1 area within each genotype, and no significance between genotypes by two-way ANOVA with Tukey’s multiple comparisons test. (**C, F, I**) Ipsilateral to contralateral ratio of % Iba-1 area for WT and KO mice indicates a modest increase in microglial density, but with no significance between genotypes by unpaired, Student’s *t*-test (*n* = 4 mice/genotype).

**Figure S2. Reactive astrogliosis induced by *ATP13A2* KO or α-synuclein PFF delivery in the amygdala and substantia nigra.** Representative images for GFAP immunostaining from amygdala (**A**) and substantia nigra (**D**) of WT and *ATP13A2* KO mice, comparing contralateral with ipsilateral PFF injection at 6 MPI. Scale bars: 500 µm. **B, E**) Digital pathology quantitation of GFAP immunostaining in **B**) amygdala and **E**) substantia nigra expressed as % area, with bars representing mean ± SEM and circles representing individual animals (*n* = 4 animals/group). *P>*0.05 by paired Student’s *t*-test comparing contralateral and ipsilateral hemispheres within each genotype, and no significance by two-way ANOVA with Tukey’s multiple comparisons test comparing between genotypes. **C, F**) Ipsilateral to contralateral ratio of % GFAP-positive area for WT and KO mice in **C**) amygdala and **F**) substantia nigra, indicates a robust increase in astrocyte density induced by PFFs, that is not significantly different between genotypes. *P>*0.05 by unpaired Student’s *t*-test.

