## Supplementary figures and images for "Neuropathology in an α-synuclein preformed fibril mouse model occurs independent of the Parkinson’s disease-linked lysosomal ATP13A2 protein"

### Supplemental Figure 1

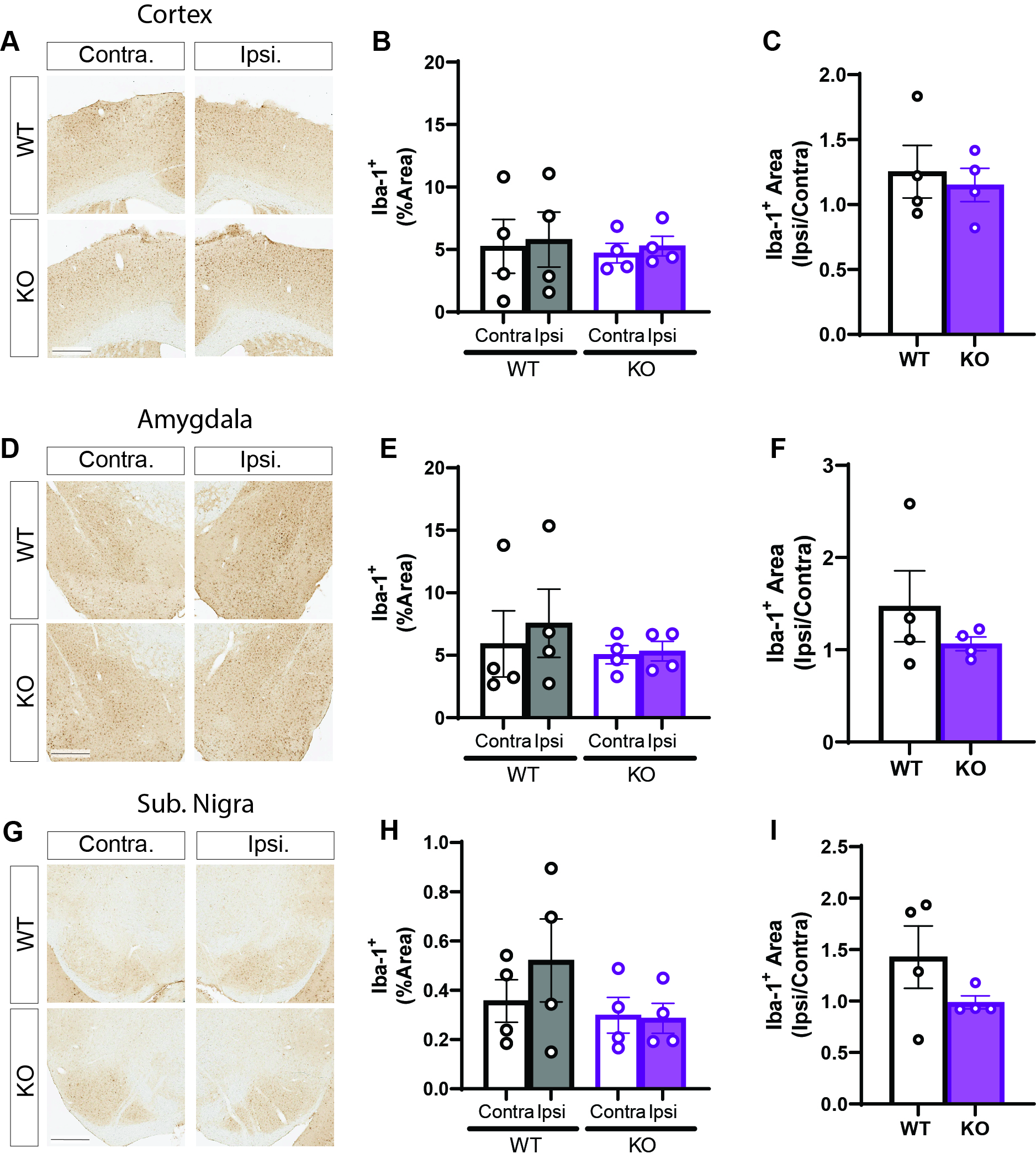

### Supplemental Figure 2

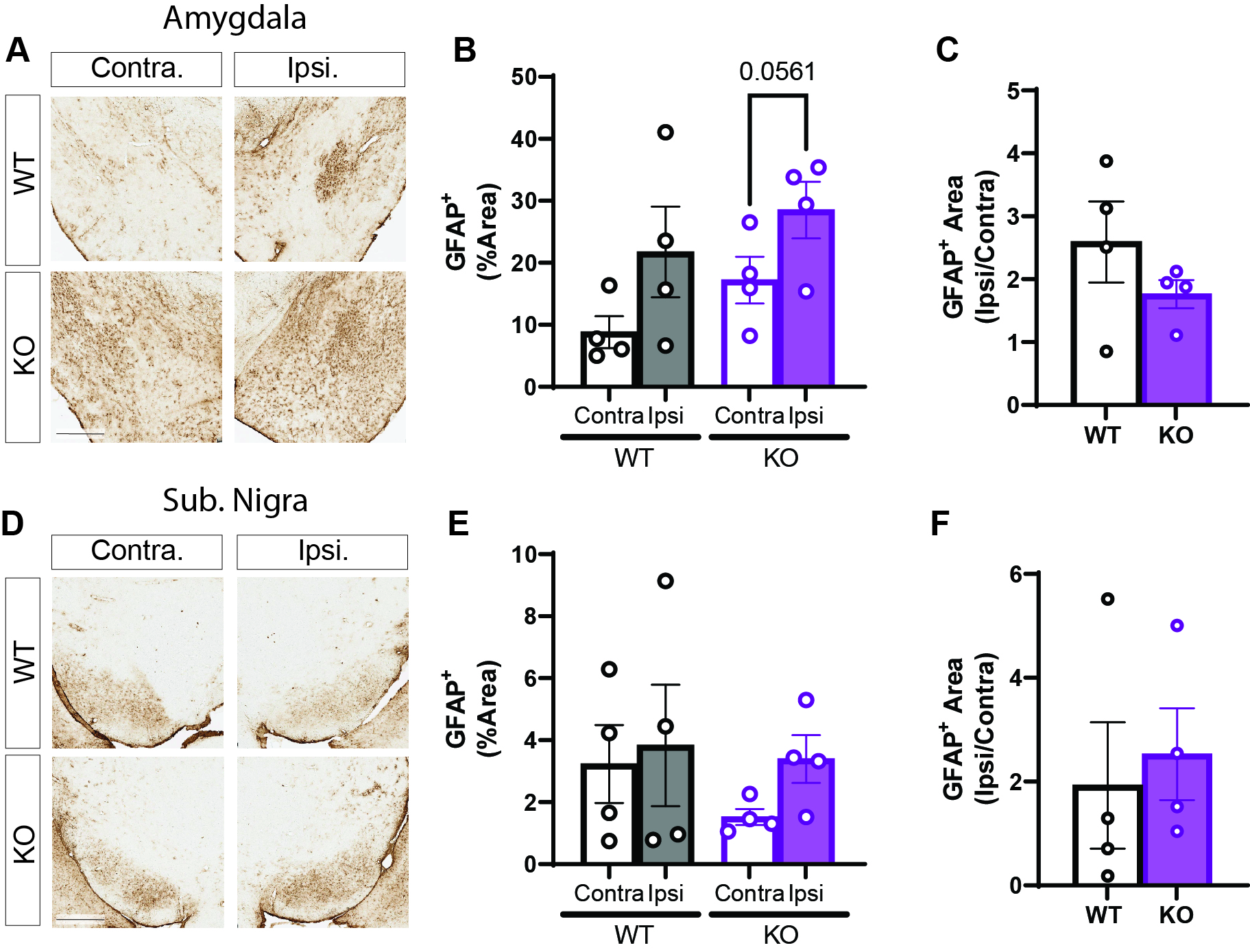
